# Homeostatic and Interferon-induced gene expression represent different states of promoter-associated transcription factor ISGF3

**DOI:** 10.1101/377275

**Authors:** Ekaterini Platanitis, Duygu Demiroz, Christophe Capelle, Anja Schneller, Markus Hartl, Thomas Gossenreiter, Mathias Müller, Maria Novatchkova, Thomas Decker

**Author notes:** These authors contributed equally (Duygu Demiroz, Christophe Capelle).

## Abstract

Cells maintain the balance between homeostasis and inflammation by adapting and integrating the activity of intracellular signalling cascades, including the JAK-STAT pathway. Our understanding how a tailored switch from homeostasis to a strong receptor-dependent response is coordinated remains limited. We used an integrated transcriptomic and proteomic approach to analyze transcription-factor binding, gene expression and in *vivo* proximity-dependent labelling of proteins in living cells under homeostatic and interferon (IFN)-induced conditions. We show that interferons (IFN) switch murine macrophages from resting-state to induced gene expression by alternating subunits of transcription factor ISGF3. Whereas preformed STAT2-IRF9 complexes control basal expression of IFN-induced genes (ISG), both type I IFN and, unexpectedly, IFNγ cause promoter binding of a complete ISGF3 complex containing STAT1, STAT2 and IRF9. In contrast to the dogmatic view of ISGF3 formation in the cytoplasm, our results suggest a model wherein the assembly of the ISGF3 complex occurs on DNA.

## Introduction

Host defense by the innate immune system requires the establishment of antimicrobial states allowing cells to cope with microorganisms before the onset of the adaptive immune response. Interferons (IFN) are of vital importance in the establishment of cell-autonomous antimicrobial immunity. Particularly the type I-IFN species IFNα and IFNβ (collectively called IFN-I) or type III IFN (IFNλ) are tightly associated with the antiviral state enabling cells to inhibit viral propagation. On the other hand, type II IFN (IFNγ), while similarly capable of inducing the antiviral state, functions predominantly as a macrophage-activating cytokine^1,2^. The accepted scenario of signal transduction by the activated IFN-I receptor complex requires the Janus kinases TYK2 and JAK1 to phosphorylate the signal transducers and activators of transcription STAT1 and 2 on tyrosine. SH2 domain-mediated heterodimerization enables STAT1-STAT2 to enter and reside in the cell nucleus. A third subunit, the interferon regulatory factor 9 (IRF9) joins the heterodimer to complete transcription factor ISGF3 which translocates to the nucleus and binds to interferon-stimulated response elements (ISREs) in ISG promoters. The IFNγ receptor on the other hand employs JAK1 and JAK2 to phosphorylate STAT1. STAT1 homodimers, a.k.a. gamma interferon-activated factor (GAF), translocate to the nucleus and stimulate ISG expression by binding to gamma interferon-activated sites (GAS)^3–5^. In addition to the canonical ISGF3 and GAF, experiments in knockout cells suggest that transcription factors containing IRF9 and either STAT1 or STAT2, but not both, have the potential to control ISG expression^6–8^. Furthermore, transcriptional activity of an ISGF3 complex assembled from unphosphorylated STATs (uSTATs) was proposed^9^. The extent to which such noncanonical complexes form and control ISG expression under conditions of a wild type cell remain elusive and are an important aspect of this study.

As the emergence of cell-autonomous immunity is an arms race between pathogen replication and restrictive mechanisms of the host, speed is a crucial attribute of the cellular response to IFN. This is particularly true for antimicrobial gene expression that must rapidly switch between resting-state and active-state transcription. Mechanisms to meet this demand include remodeling and modification of promoter chromatin prior to the IFN response^10^. Moreover, a host of studies support the concept that cells permanently produce a small amount of IFN-I that stimulates a low, tonic signal by the IFN-I receptor^11,12^. This in turn generates a baseline transcriptional response of IFN-induced genes (ISG). In support of this idea, disrupting IFN-I production or signalling in resting-state cells was shown to cause a drop in basal ISG expression. The enhancement of ISG transcription by an IFN stimulus from basal to induced levels has been compared with the revving-up of a running engine^13^. The ‘revving-up’ or ‘autocrine loop’ model predicts a tight coupling between homeostatic and receptor mediated interferon signalling and that transcriptional ISG activation in resting and activated states differs in its intensity, but abides by the same mechanism.

Our study challenges this notion. By combining ChIP-seq and transcriptome analysis we find that basal expression of many ISGs is controlled by a preformed STAT2-IRF9 complex, whose formation does not require signalling by the IFN-I receptor. IFN treatment induces a rapid switch from the STAT2-IRF9 to the canonical ISGF3 complex, revving up ISG transcription. Quantitative proteomic analysis and in vitro interaction studies suggest a model wherein the majority of IRF9 resides in the nucleus under homeostatic conditions and the assembly of the ISGF3 complex occurs on DNA. In conclusion, combining high-throughput data enabled us to reveal mechanisms by which different states of promoter-associated transcription factor ISGF3 control the switch from homeostatic to interferon induced gene expression.

## Results

### Different transcription factors bind to ISG promoters under resting and induced conditions

We used macrophages to study mechanisms contributing to constitutive ISG expression. Basal expression of ISGs controlled via ISRE promoter sequences (Irf7, Usp18 and Oas1a) in bone marrow-derived macrophages (BMDM) was strongly reduced by ablation of any of the three ISGF3 subunits (Fig. 1a). Surprisingly, genes regulated predominantly by STAT1 dimer binding to GAS sequences such as Irf1 and Irf8 were largely unaffected by the gene deficiencies including STAT1 (Fig. 1b). Since IRF9 is the DNA binding subunit of all ISRE-associated transcription factors, our results point towards an important role of this protein and its associates for basal ISG expression. Consistent with this, RNA-Seq and gene set enrichment analysis (GSEA; Fig. 1c) underscore the impact of IRF9 loss on global ISG transcription in resting cells. To determine whether IRF9 dependence reflected the formation of an ISGF3 complex we performed ChIP-Seq in wt cells. The integrated experimental approach is shown in supplementary Fig.1. This first-ever examination of all three ISGF3 subunits simultaneously was made possible by the generation of a novel anti-IRF9 monoclonal antibody, 6FI-H5, which yields excellent signal-to-noise ratios in ChIP (Supplementary Fig. 2a). IRF9-dependence of STAT1/2 binding was confirmed by ChIP-Seq in Irf9-/- BMDM.

**Figure 1.**
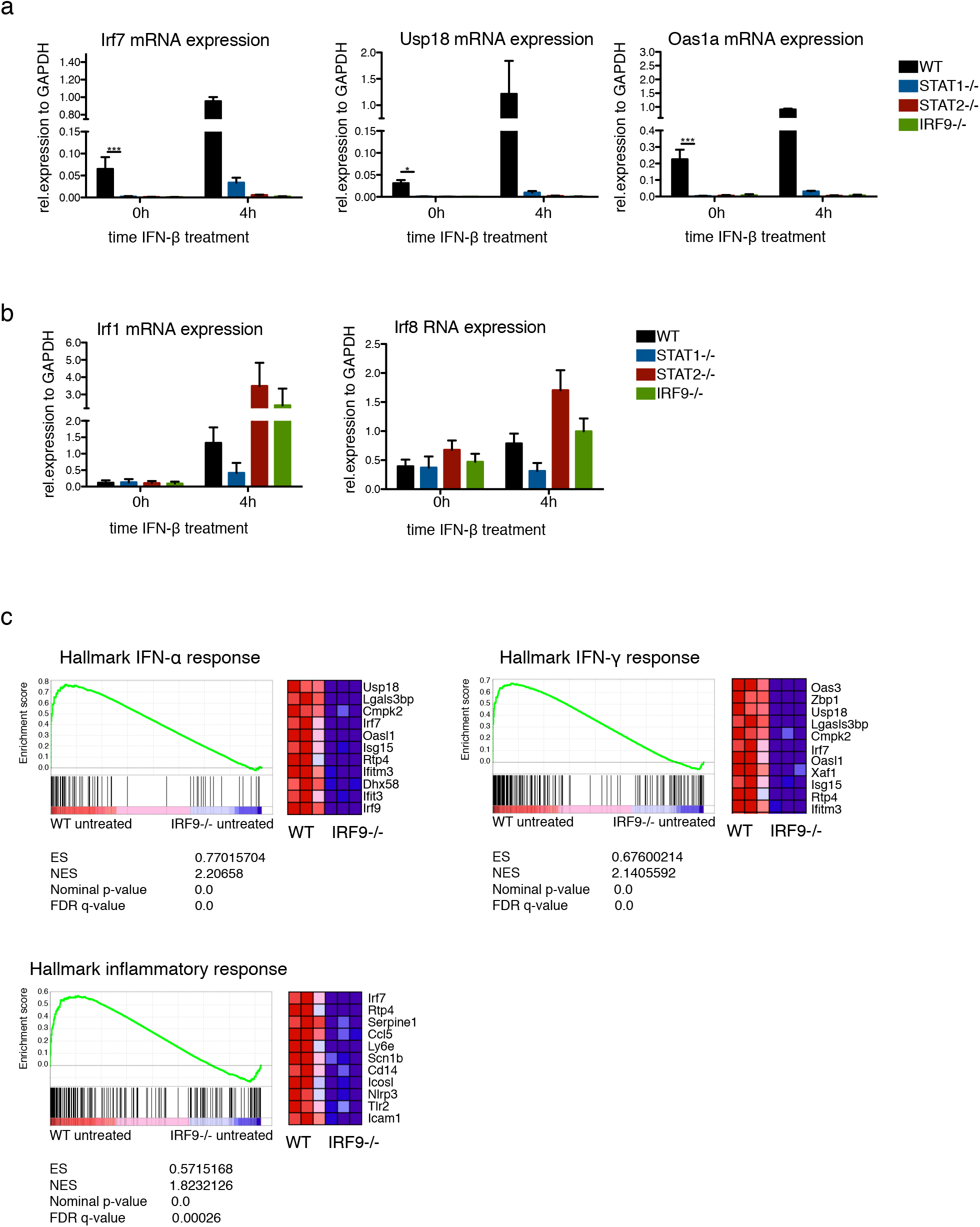
Conditions of basal ISG expression. a, b) Bone marrow-derived macrophages (BMDM) isolated from wild-type (WT), Stat1^-/-^, Stat2^-/-^, and Irf9^-/-^ mice were treated with 250 IU/ml of IFN-β as indicated. Gapdh-normalized gene expression was measured by RT-qPCR. Data represent the mean and standard error of the mean (SEM) values of three independent experiments. P-values were calculated using the paired ratio t-test (\**P* ≤ 0.05; \*\**P* ≤ 0.01, \*\*\**P* ≤ 0.001). c) Gene set enrichment analyses showing upregulation of an IFN and inflammatory response signature of untreated WT compared to untreated Irf9-/- BMDM. The top correlated genes for each biological triplicate are displayed in the corresponding heat maps. The total height of the curve indicates the extent of enrichment (ES), with the normalized enrichment score (NES), the false discovery rate (FDR) and the *P* value.

The combination of RNA-Seq (plotting wt versus Irf9-/- cells) and ChIP-Seq data (revealing promoter occupancy in wt cells) in a scatter plot showed promoters of a majority of IRF9-dependent genes associated with STAT2 and IRF9, but not with STAT1 (Fig. 2a, d. A quantitative representation of promoters binding STAT2-IRF9, ISGF3 or STAT1 dimers is given in the pie chart inserts and genes are listen in supplementary table 1). A much smaller fraction was associated with all subunits of an ISGF3 complex. A brief treatment with either IFN-I or IFNγ caused a vast majority of promoters, including many of those associated with STAT2-IRF9 in resting state, to bind ISGF3 (Fig. 2b,c and 2e,f). Promoters associated with STAT1 dimers were prominently represented among IFNγ-induced genes, but not among IFN-I-induced ISG. According to the prevailing JAK-STAT paradigm transcriptional IFN-I responses are ISGF3/ISRE based while those to IFNγ use STAT1 dimers/GAS. Surprisingly we observed a prominent contribution of ISGF3 to IFNγ-induced ISGs as well as the de novo formation of STAT2-IRF9 complexes at both IFN-I and IFNγ-induced ISGs. Among the overlapping peak sets a small number of IRF9 complexes can be found which, upon visual inspection, appeared to be co-bound by STAT2 and are therefore more likely the result of type II peak prediction error (Supplementary Fig. 2b). Taken together the data are consistent with the notion that the switch from resting-to activated-state transcription is caused by a transition from STAT2-IRF9 to ISGF3 at a majority of ISG promoters.

**Figure 2.**
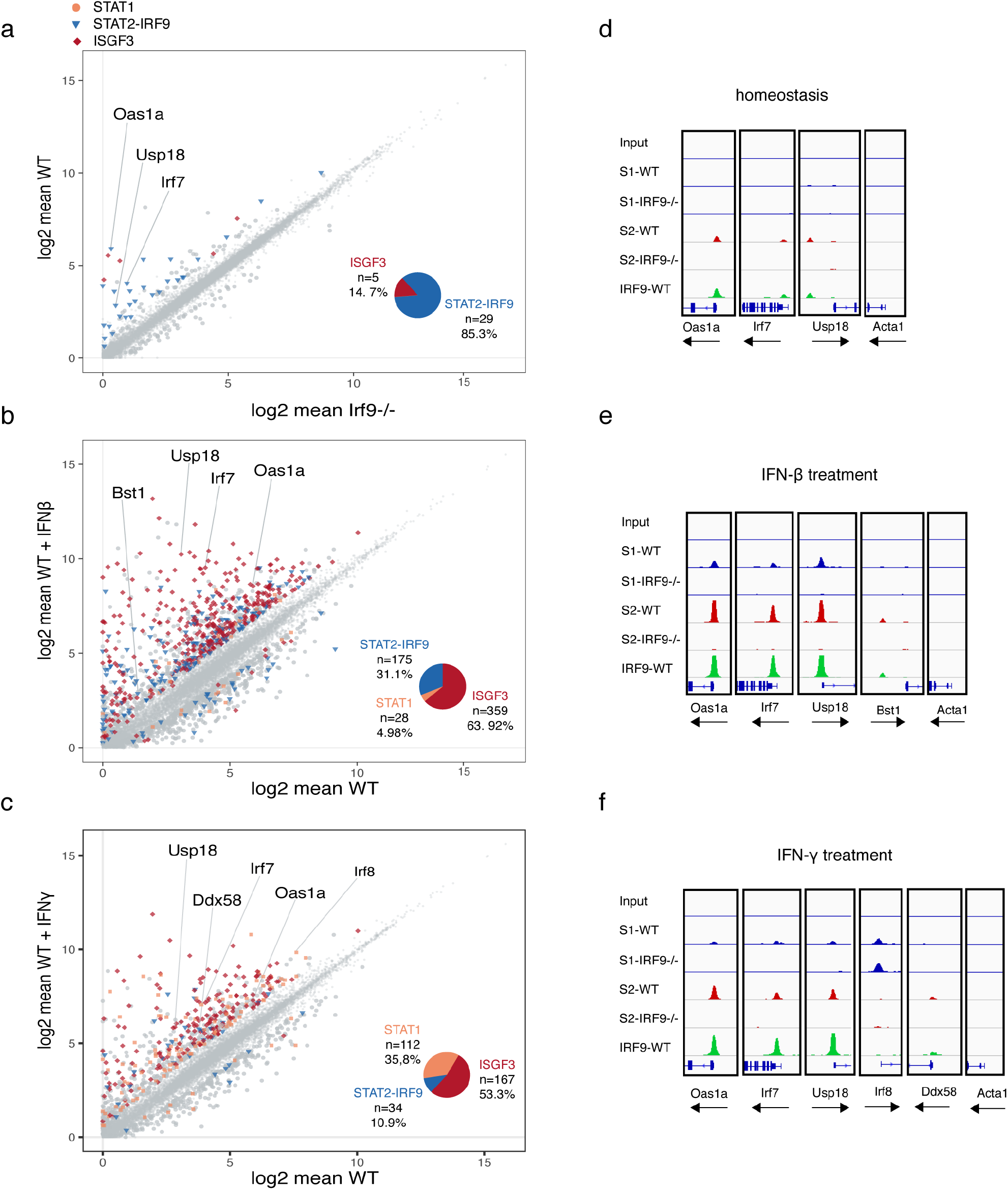
IFN-induced changes in gene expression and chromatin-associated STAT complexes. a-c) Scatterplot linking RNA-seq (n=3) with ChIP-seq (n=2) data. Significantly differentially expressed genes between IRF9-/- and WT, WT untreated and WT IFN-β as well as WT untreated and WT IFNγ are displayed in color for abs (logFC)>1. ChIP-seq analysis was performed on untreated (a), IFN-β (b) or IFNγ (c) treated BMDMs of wild-type (WT) and Irf9-/- (IRF9-/-) mice. Two independent biological replicates were used for the analysis. Chromatin immunoprecipitation (ChIP) was performed using STAT1, STAT2 and IRF9 antibodies. Symbols indicate bound complexes as determined by peak coincidence. d-f) Genomic analysis of STAT1 (S1) (scale 0-150), STAT2 (S2) (scale 0-200) and IRF9 (scale 0-200) binding. Transcription factor binding at ISG and control loci (scale 0-200). See also Supplementary Figure 1 and 2.

### Complex formation and intracellular proximity of ISGF3 subunits

Co-immunoprecipitation studies in human epithelial cell lysates concluded that STAT2 and IRF9 form complexes devoid of STAT1. Corroborating this result, recent work solved the structure of the binding interface and determined that STAT2 binds IRF9 with 500-fold higher affinity than STAT1^14^. Besides, STAT2-IRF9 as well as STAT1-STAT2 complexes could be co-immunoprecipitated from resting-cell extracts^15–18^. Our results in mouse BMDM are in agreement with these observations. Reciprocal immunoprecipitations demonstrated a clear association between STAT2 and IRF9 that increased after IFN-I treatment (Fig. 3a). Furthermore, we observed a weak association between STAT2 and STAT1 that, in line with earlier observations, did not increase after IFN-I treatment^15,17^. STAT1 and IRF9 could not be coprecipitated despite earlier studies assigning transcriptional activity to STAT1-IRF9 complexes^19,20^. Thus, while STAT2-IRF9 and a significantly smaller amount of STAT1-STAT2 complexes can be demonstrated in resting cells, we observed neither ISGF3 nor other complexes containing both STAT1 and IRF9. To corroborate these findings and to rule out that the data reflected stability under IP conditions rather than complex formation in cells, we used the BioID proximity labelling technology as depicted in Supplementary Fig. 1. Modified biotin ligase BirA* fusion proteins^21^ biotinylate proteins within a distance of approximately 10nm^22^. Raw 264.7 macrophages, which show the same response to IFN as BMDM (Supplementary Fig. 3a), were engineered to express a doxycycline (Dox)-inducible, myc-tagged IRF9-BirA* fusion gene and used to study complex formation in vivo. The fusion protein restored IFN signalling in Irf9-/- mouse embryonic fibroblasts (Supplementary Fig. 3b). To avoid overexpression artefacts, we matched IRF9-BirA* with endogenous IRF9 levels (Supplementary Fig. 3c) and used cells expressing an N-terminally myc tagged BirA* gene as controls. We examined STAT-IRF9 proximity prior to IFN treatment by parallel reaction monitoring (PRM), an approach allowing to specifically acquire information about peptides of interest and their quantities^23^. PRM revealed that IRF9-BirA* biotinylated both itself and STAT2 in two biological replicates (Fig. 4a, Supplementary table 2). In contrast, STAT1 was not enriched by Streptavidin-mediated affinity purification compared to the control cells. A 90 min pulse with either IFN-I or with IFNγ did not cause detectable STAT1-IRF9 proximity (Fig. 4b) although it caused ISGF3 formation on DNA (Fig. 2). STAT2 on the other hand was among the top interactors under these conditions (Fig. 4b; Supplementary Fig. 4a). Further analysis of the constitutive IRF9 interactome revealed an association between IRF9 and a number of chromatin modifiers (Fig. 4b-d; Supplementary Fig. 4a).

**Figure 3.**
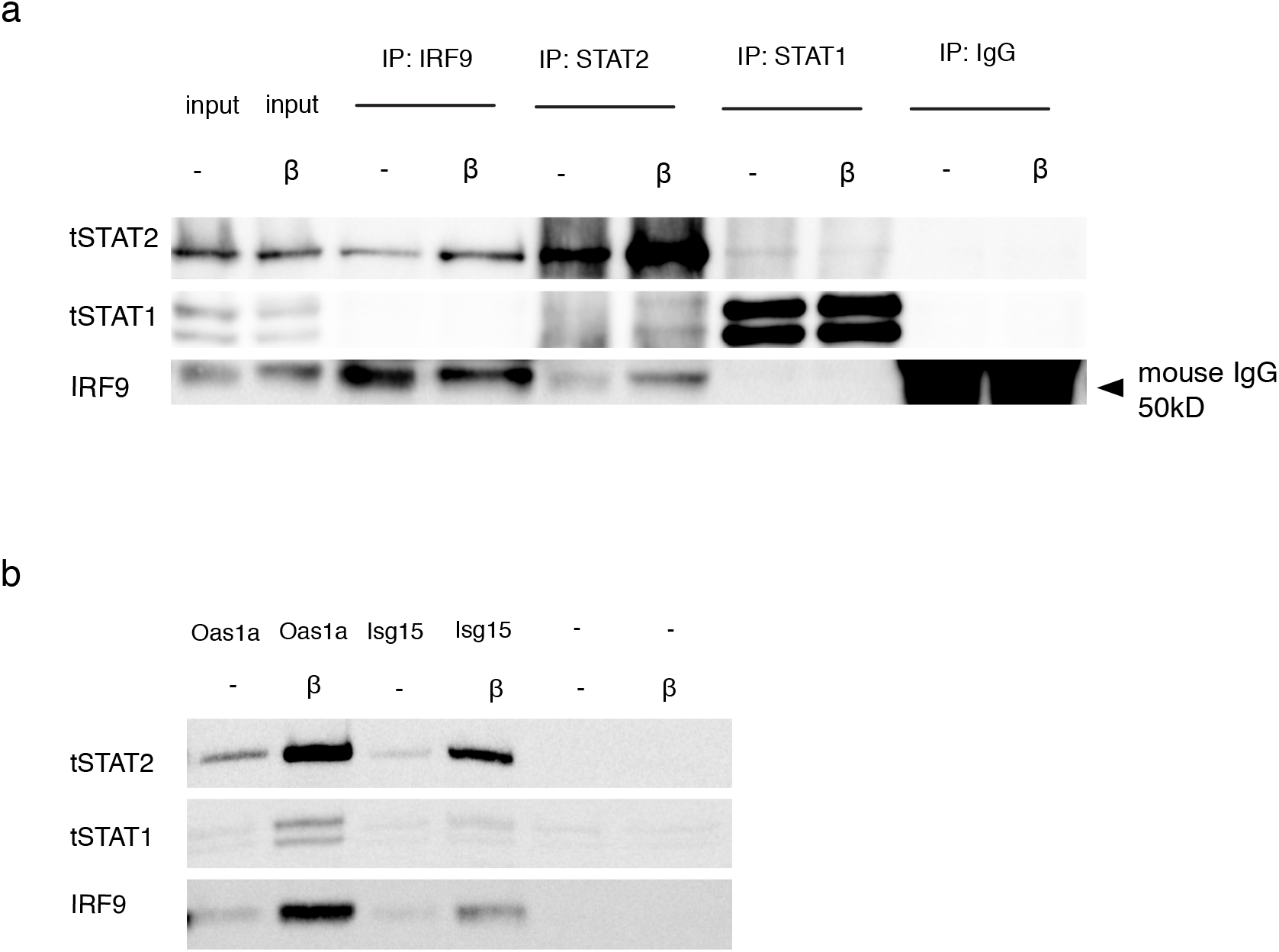
Signal- and DNA dependence of complex formation from ISGF3 subunits. a) BMDMs from wild-type (WT) animals were treated for 1.5h with IFN-β and immunoprecipitation was carried out using antibodies against STAT1, STAT2, IRF9 or an IgG control. Immunoprecipitated complexes were analyzed by western blotting with antibodies to STAT1, STAT2 and IRF9. Input controls represent 10% of the total lysate used for the immunoprecipitation. b) Raw 264.7 cells were treated for 1.5h with IFN-β. Lysates were either incubated with a biotinylated oligo harboring the Oas1a-ISRE or the Isg15-ISRE sequence, or plasmid DNA. DNA bound protein complexes were isolated by streptavidin affinity purification, followed by western blot analysis.

**Figure 4.**
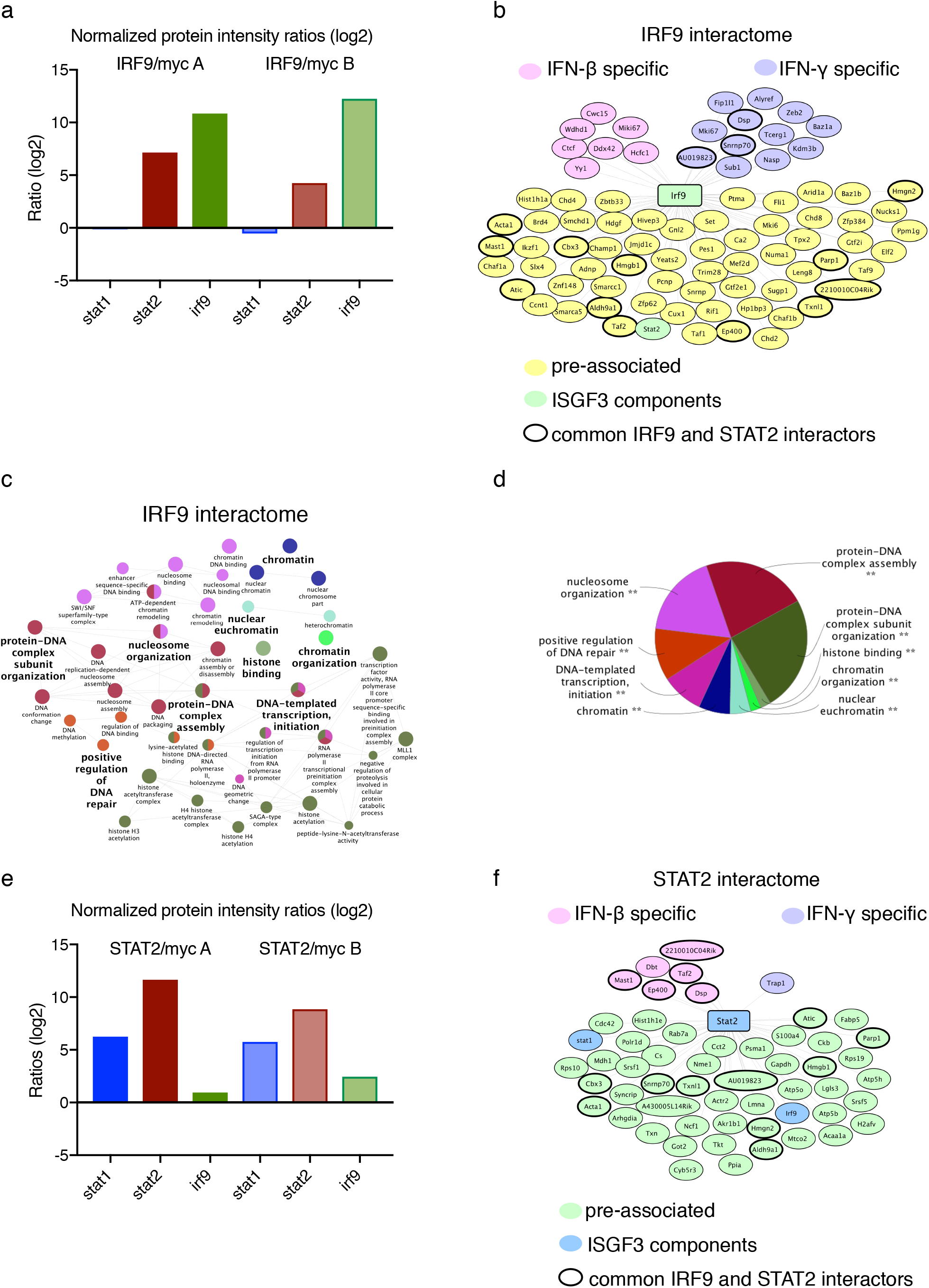
Affinity labeling of IRF9 and STAT2 interactors. a), e) Analysis of IRF9 (a) and STAT2 (e) BioID PRM runs using Skyline software. RAW264.7 cells were treated with 0,2μg/ml doxycycline for 24h, followed by addition of 50μM biotin for 18 hours. Cells were lysed and protein complexes isolated by streptavidin affinity purification, followed by analysis with LC-MS. Log2-transformed protein ratios were calculated for two biological replicates from mycIRF9-BirA/mycBirA cells (a) or mycSTAT2-BirA/mycBirA cells (e). b), f) Network diagram of IRF9 (b) or STAT2 interactors (f), generated with Cytoscape software. Proteins that displayed a threefold enrichment in the experimental samples compared to the ligase only controls were counted as hits and are displayed in the network. Spatial arrangement of nodes and group color coding reflects proteins either pre-associated in the absence of any signal or specifically enriched upon 1.5h of IFN-β or IFN-γ treatment. Distance of connecting lines (edges) between proteins do not reflect the strength of the interaction. c) ClueGO network analysis of the IRF9 interactome. The app was used to find overrepresented GO processes and a network of connected GO terms was created. Each node represents a GO biological process, and the colors represent the GO group. d) Pie chart of ClueGO results. See also Supplementary Figure 1, 3 and 4.

To rule out that the lack of interaction between IRF9 and STAT1 resulted from the unavailability of lysines for biotinylation, we generated Raw 264.7 macrophages expressing a Dox-inducible, myc-tagged STAT2-BirA* fusion gene. STAT2-BirA* expressed at endogenous levels was functional and restored the IFN response of Stat2-/- MEFs (Supplementary Fig. 3d, e). PRM confirmed IRF9 proximity to STAT2-BirA* and additionally revealed proximity between STAT2 and STAT1 (Fig. 4e). The latter finding was further verified in western blots of STAT2-BirA*cell lysates with HRP-coupled streptavidin (Supplementary Fig. 3e). Treatment with IFN further increased the interaction between STAT2 and IRF9 (Supplementary Fig. 4b). Importantly, the spectrum of STAT2 interactors partially overlapped with that produced in the IRF9-BioID approach (Fig. 4b, f, supplementary table 2).

Together, the co-IP experiments and the BioID approach demonstrate the presence of STAT2-IRF9 and of STAT1-STAT2 heterodimers in resting cells and confirm our finding that neither resting conditions nor IFN-treatment produce detectable amounts of complexes containing both STAT1 and IRF9. This suggests that ISGF3 complexes may not form in absence of DNA. To examine this possibility, we carried out DNA-mediated precipitation studies (Fig. 3b). Oligonucleotides representing ISG15 or Oas1a ISREs precipitated STAT2-IRF9 from resting-cell extracts, but the entire ISGF3 complex after stimulation with IFN-I (the weak STAT1 band in resting cell extracts (lane 1) was also observed in absence of DNA (lanes 5 and 6) and results from nonspecific association to the carrier material). This behaviour resembles complex binding in ChIP-Seq (Fig. 2). We conclude that ISRE binding either stabilizes ISGF3 or is required for its formation.

### Signalling requirements for STAT2-IRF9 formation

The revving-up model of innate immunity predicts that the molecular machinery for constitutive ISG expression requires a low chronic signal from the IFN receptor. It further implies that a small quantity of ISGF3 subunits should be nuclear in untreated cells. To test these assumptions, we first analysed the cellular localization of ISGF3 subunits. We have recently shown that IFN-induced nuclear localization of STAT2 is reduced in Irf9-/- BMDMs^24^. It follows that a portion of IRF9 should be localized to the cell nucleus and that nuclear shuttling of at least a sub-fraction of STAT2 requires IRF9. Indeed, the majority of IRF9 in resting cells was found in the nucleus by immunofluorescence and IFN-I treatment did not further enhance nuclear accumulation (Fig. 5a). Raw 264.7 cell fractionation and western blots revealed that nuclei from resting cells contained IRF9 and a small fraction of STAT2, but not STAT1 (Fig. 5b). The small quantity of STAT1 bound to chromatin according to ChIP-Seq appears to be below the detection limit. To assess whether IFN receptor signalling was important for STAT2 nuclear accumulation, we treated cells with Staurosporine, a potent kinase inhibitor, prior to IFN stimulation. The inhibitor did not reduce nuclear STAT2 amounts in resting cells. By contrast, IFN-induced nuclear translocation of STAT2 was abrogated. The small STAT2 quantity found in the nucleus of IFN-I and inhibitor treated cells equaled that of unstimulated cells (Fig. 5b). Next, the cells were treated with a specific JAK inhibitor, P6, which completely abrogated STAT1 and STAT2 tyrosine phosphorylation in IFN-I and IFNγ treated BMDMs (Fig. 5c). As anticipated, the addition of P6 prior to a short IFN treatment led to a complete loss of STAT1 phosphorylation at Y701 and inhibition of STAT1 translocation to the nucleus (Fig. 5d). In contrast, a small fraction of STAT2 was detected in the nucleus of both Raw 264.7 macrophages (Fig. 5e) and BMDM (Fig. 5f), despite a complete loss of Y689 phosphorylation. Taken together these observations support the conclusion that the formation and nuclear localization of STAT2-IRF9 complexes occur independently of a continuous signal from the IFN-I receptor.

**Figure 5.**
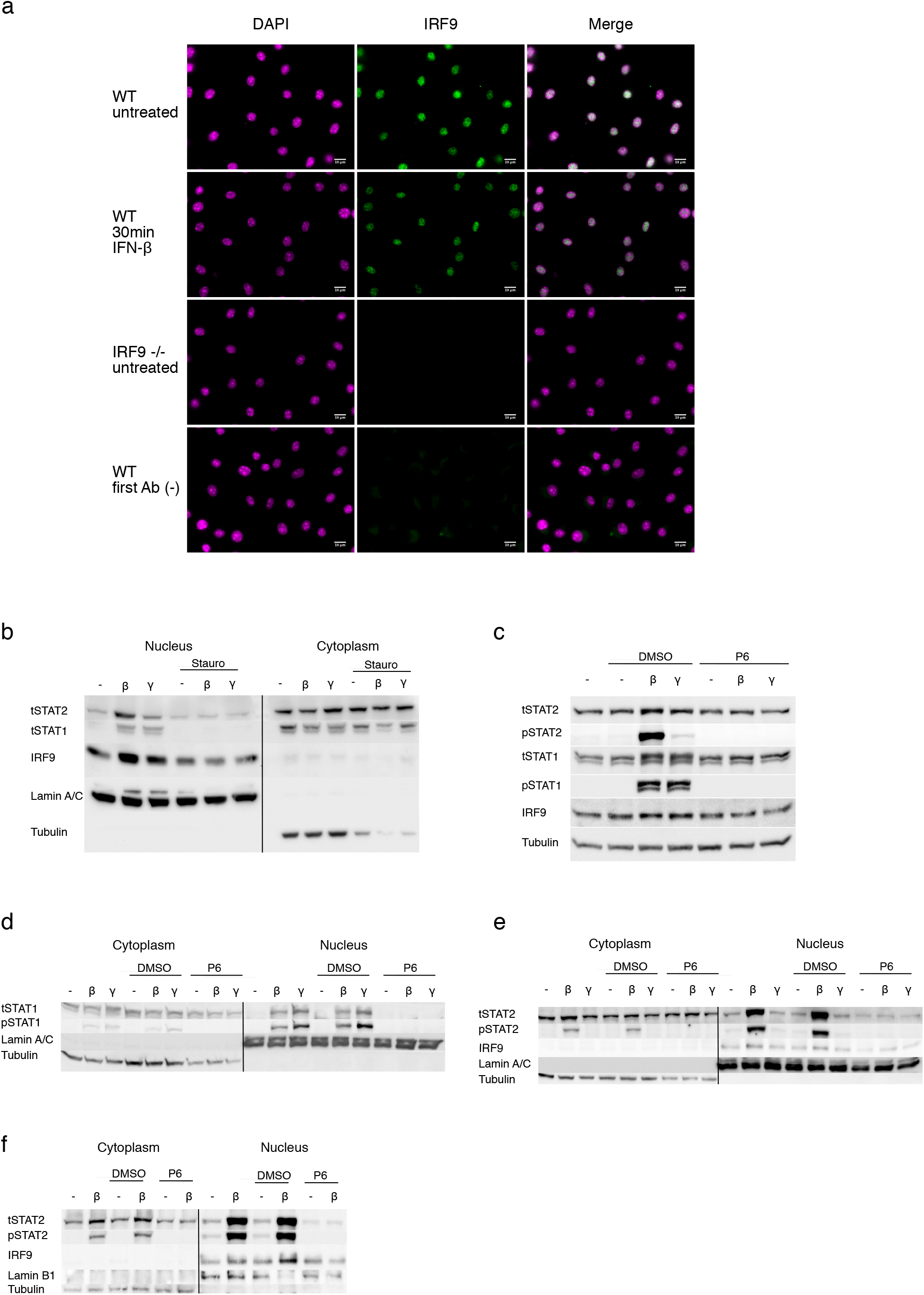
Localization of STAT complexes. a) IRF9 localization as determined by immunofluorescence. BMDMs of wild-type (WT) and Irf9-/- (IRF9-/-) mice were left untreated or stimulated with 250 IU/ml of IFNβ for 30 min. The cells were fixed and stained with anti-IRF9 antibody followed by Alexa Fluor^®^ 488 conjugated secondary antibody (green). Nuclei were stained with DAPI (magenta). First Ab (-) indicates control without the first antibody. The scale bars represent 10 μm. b,d, e) Raw 264.7 nuclear and cytoplasmic extracts were prepared from controls or after a 30-minute treatment with IFN-β or IFN-γ and analyzed by western blot. The entire nuclear fraction and 30% of the cytoplasmic fraction are shown. b) STAT1, STAT2, IRF9 and α-Tubulin, Lamin A/C controls are shown. 500nM staurosporine was added 3h prior to IFN treatment. c) Phosphorylation of STAT1 at Y701, STAT2 at Y689, total STAT1, total STAT2, IRF9 and α-Tubulin of total bone marrow derived macrophage lysates were determined. 15μM P6 inhibitor or DMSO were added for 3h prior to IFN treatment d) Phosphorylation of STAT1 at Y701, total STAT1, α-Tubulin and Lamin A/C levels were determined. 15μM P6 inhibitor or DMSO were added for 3h prior to IFN treatment. e) Phosphorylation of STAT2 at Y689, total STAT2, IRF9, α-Tubulin and Lamin A/C. 15μM P6 inhibitor or DMSO were added for 3h prior to IFN treatment. f) Nuclear and cytoplasmic extracts from control wt-BMDM or after 30-minute treatment with IFNβ were analyzed for the phosphorylation of STAT2 at Y689 as well as total STAT2, IRF9, α-Tubulin and Lamin B1. 15μM P6 inhibitor or DMSO were added for 3h prior to IFN treatment.

### Tonic IFN receptor signalling maintains expression of ISGF3 subunits

The continued presence of STAT2-IRF9 in the nucleus suggests that basal ISG expression should be largely unaffected by JAK inhibition. To test this hypothesis, untreated or P6-treated WT BMDM were compared to the ISGF3-subunit knock-outs (knockout data are as in Fig. 1 and included here for ease of comparison). In agreement with our assumption, basal expression of genes with pre-bound STAT2-IRF9 was largely maintained in the presence of inhibitor (Irf7, Usp18, Oas1a; Fig. 6a). All three knockouts strongly affected STAT2-IRF9-dependent genes. Genes induced via GAS sequences (Irf1, Irf8) were unaffected by either the inhibitor or the knockouts. Thus, shutting down signalling while maintaining ISGF3 subunit levels, as is the case in our inhibitor experiments, sustains basal ISG expression independent of tonic IFNAR signalling.

**Figure 6.**
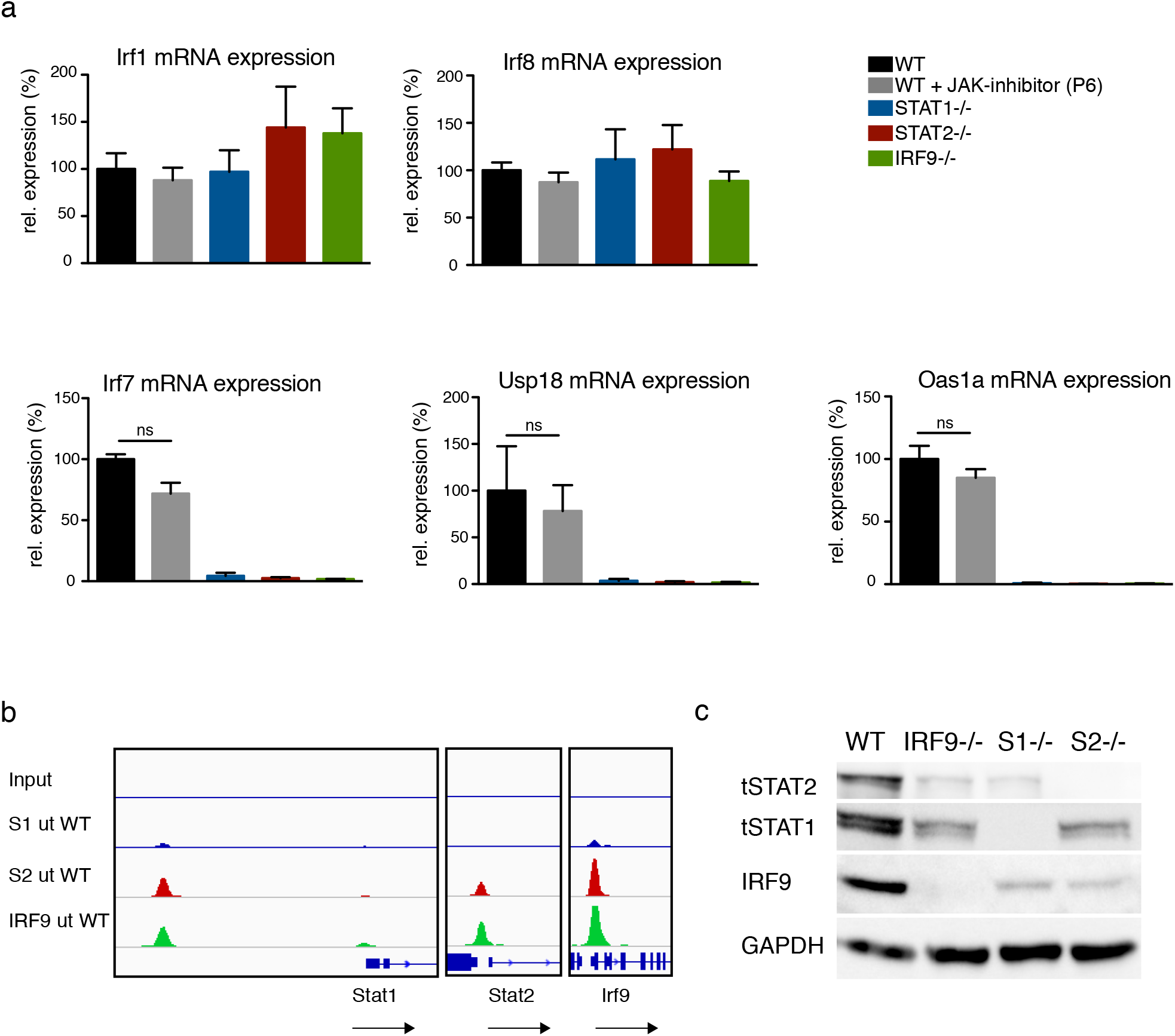
Cross regulation of ISGF3 subunits. a) BMDMs isolated from WT, Stat1^-/-^, Stat2^-/-^, and Irf9^-/-^ mice were left untreated or treated with 15μM P6 inhibitor for 3h. Gapdh-normalized gene expression was measured by RT-qPCR. Data represent relative expression in percentage, where WT untreated equals 100%. Data represent the mean and standard error of the mean (SEM) values of three independent experiments. *P*-values were calculated using the paired ratio *t*-test (\**P*≤ 0.05; \*\**P* ≤ 0.01, \*\*\**P* ≤ 0.001). b) Genomic analysis of STAT1 (S1), STAT2 (S2) and IRF9 binding. Transcription factor binding at Stat1, Stat2 and Irf9 transcriptional start site. Input, STAT2 and IRF9 (scale 0-200). STAT1 (scale 0-150) c) Whole-cell extracts from wild-type, Stat1^-/-^, Stat2^-/-^ and Irf9^-/-^ BMDMs were tested by western blot for total STAT1, STAT2 and IRF9 levels (representative of three independent experiments).

The persistence of nuclear STAT2-IRF9 upon JAK inhibition raises the question of why genes whose basal expression is sustained by this complex are sensitive to a permanent disruption of signalling in cells lacking the IFN-I receptor or STAT1, or that express a STAT1Y701F mutant^24^. A straightforward answer to this question is provided by the results in Fig. 6b showing that ISGF3 subunits are bound to the promoters of each of their genes in resting cells and suggesting they all contribute to each other’s basal expression. Consistent with this, Stat1 and Irf9 promoters were associated with ISGF3 and the Stat2 promoter bound STAT2-IRF9. Thus, gene deletion of all three ISGF3 subunits is expected to lower IRF9 levels and therefore any ISRE-dependent basal expression. Consistently, knockout of each subunit caused a severe reduction of the two other subunits (Fig. 6c). Thus, cells lacking the ability to form an ISGF3 complex express low amounts of all its subunits and are therefore unable to sustain STAT2-IRF9-dependent basal gene expression. The data emphasize the importance of studying STAT complex formation in wt conditions. An integrated model depicting the proposed interplay between tonic signalling-dependent and independent events for basal ISG expression and the changes occurring upon IFN treatment are depicted in Fig. 7.

**Figure 7.**
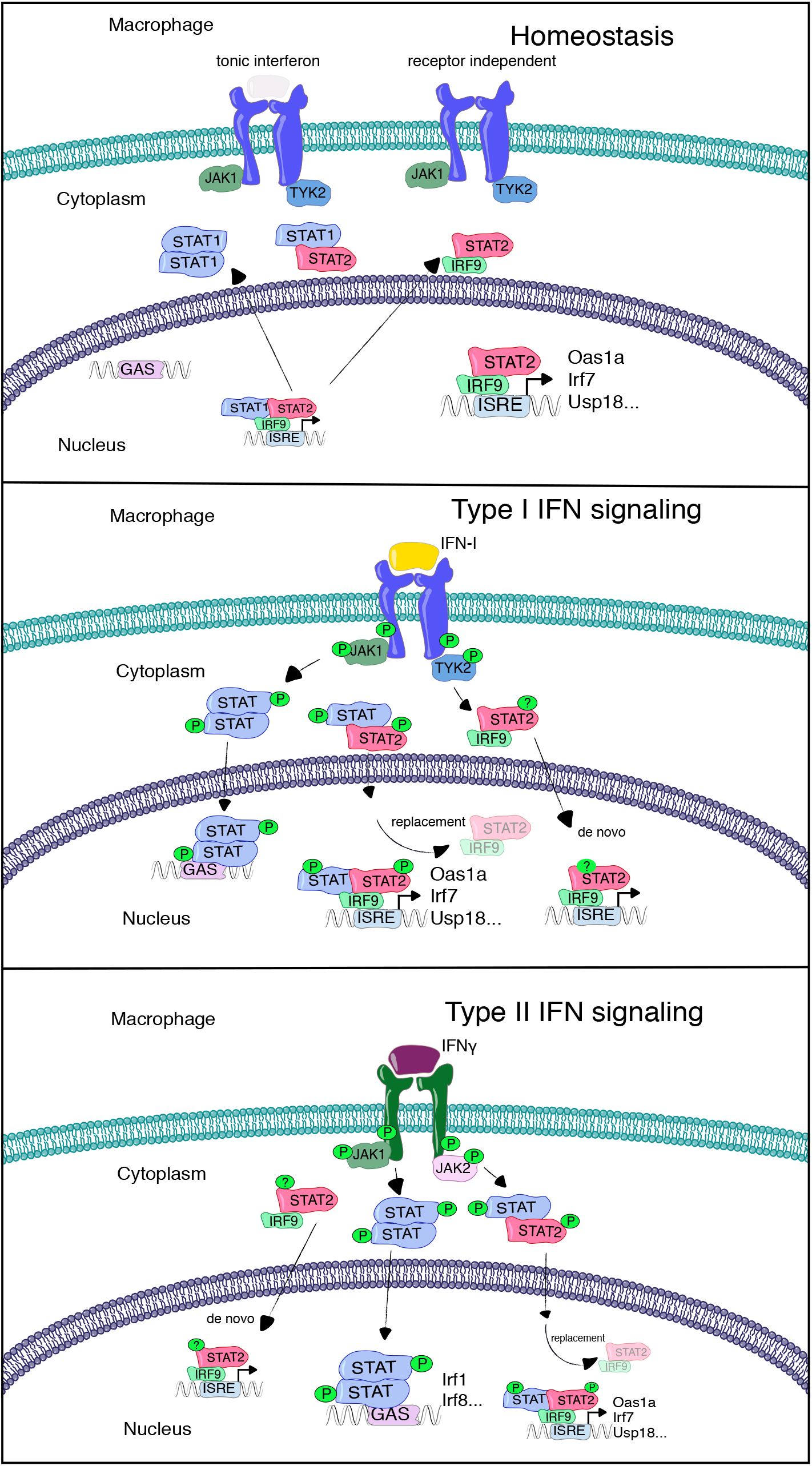
Model depicting the molecular switch from resting to IFN-induced gene expression. Whereas a preformed, signalling-independent STAT2-IRF9 complex drives the basal expression of interferon-induced genes, both IFN-I and IFN-γ cause increased transcription by replacing this complex with ISGF3. For detailed information see text.

## Discussion

Recent findings suggest that unphosphorylated STAT complexes control ISG expression^8^. Our study shows an important contribution of STAT2-IRF9 complexes, formed independently of IFN-I receptor signalling, to constitutive ISG expression in resting cells. This added mechanistic insight calls for a reinterpretation of the ‘revving-up’ model and the implications of tonic signalling by the IFN-I receptor^12,13^. Both models provide a mechanistic framework to explain how cells are prepared for the race between pathogen multiplication and innate resistance. They posit that IFN-dependent responses such as the antiviral state or macrophage activation are not established de novo during immune responses, but represent the enhancement of a pre-existing condition compatible with resting-state physiology. We confirm that tonic signalling is indeed of critical importance for the maintenance of ISGF3 subunits and show that the signal-independent formation of STAT2-IRF9 complexes, previously considered as non-canonical, is an integral component of ISG regulation. The change from STAT2-IRF9 to ISGF3 functions is a transcriptional mode switch between resting and active states for many ISGs. Importantly, our observations rely on wt cells. The data with knockout cells show that deletion of one STAT subunit not only extinguishes the function of that protein, but also changes the stoichiometry of the remaining ISGF3 components. Component stoichiometry is an important determinant for the proper biological output of JAK-STAT signalling. For example, IL-6 signalling in the absence of STAT3 exhibits a transcriptional profile similar to that of IFNγ by preferentially using STAT1^25^.

Basal ISG expression requires the constant secretion of miniscule IFN-I amounts to prime cells for a rapid response to microbial infections. This is illustrated by IFNβ promoter-luciferase reporter mice that show weak luminescence in the entire body^26^. Earlier studies revealed IFN-I mRNA and protein in tissues under pathogen-free conditions^27,28^. Besides priming cells for enhanced cytokine responsiveness, this homeostatic mechanism reveals its importance in the maintenance and mobilization of the hematopoietic stem cell niche^29,30^. More recent studies suggest that multipotent stem cells achieve ISG expression and an antiviral state independently of IFNAR signalling^31^. The results strengthen the idea that mobilization of ISG promoters via STATs can be uncoupled from IFN-I receptor signalling. The uSTAT model provided a first proof of principle for this possibility. Accordingly, the prolongation of ISG expression occurs via formation of a uISGF3 complex following an early, IFNAR-dependent response that increases the levels of ISGF3 subunits^9^. In hepatic cells a uISGF3 was proposed to function under homeostatic conditions and without a previous IFNAR-dependent signal^32^. However, a mechanism explaining IFNAR-independent ISG expression under conditions of a wildtype cell remained elusive. Here we show that the switch of many ISGs from basal to rapid IFN-induced expression requires exchanging STAT2-IRF9 for ISGF3 complexes. Our model receives strong support from a study comparing basal gene expression in Ifnar1-/- and Tyk2-/- cells, both lacking tonic signalling^33^. This revealed a large overlap with genes whose expression was impaired by the loss of IRF9 in our study. Importantly, tonic-sensitive loci showed higher STAT2 binding at baseline^33,34^. Strong support for a role of IRF9-STAT2 in basal gene expression is also provided by the finding that an IRF9 fusion protein with the STAT2 transactivating domain is sufficient to increase basal ISG expression and to induce an antiviral response in absence of added IFN^35^.

Earlier studies showing pre-association of STAT2 and IRF9 proposed cytoplasmic localization of the complex, due to a dominant STAT2 nuclear export sequence^36^. Our data revise this idea by showing that a proportion of STAT2-IRF9 resides in the nucleus of resting cells. Importantly, our antibody- and DNA-mediated co-precipitation studies argue in favor of an ISGF3 formation on DNA. In support of this, earlier biochemical studies have failed to demonstrate a DNA-independent, complete ISGF3 complex^37^. More generally, the switch from STAT2-IRF9 to ISGF3 poses the question how these complexes manage constitutive and IFN-induced ISG expression. One reason for the transcriptional increase associated with ISGF3 binding is most likely the improved stability on DNA. While IRF9 makes contacts to the core ISRE, STAT1 increases ISGF3 affinity by additionally interacting with 5’ flanking sequences^38^. A higher off-rate of STAT2-IRF9 would also explain how this transcription factor can be rapidly exchanged for ISGF3 after IFN treatment. An additional implication might be transcription factor cooperativity. For example, a STAT2-IRF9 complex reportedly allows for co-regulation of the human IL6 gene by IFN and NFκB-activating agents because STAT2 binds IRF9 as well as NFκB p65^39^. Consistently, data by Mariani et al. suggest that STAT2-IRF9 mediates cooperativity required for the enhanced induction of genes in response to IFNβ and the NFκB-activating cytokine TNF^40^. Thus, it is possible that STAT2-IRF9 and ISGF3 represent alternative platforms for the integration of transcription factor signalling at promoter level.

In vivo biotinylation shows IRF9 to be in close proximity to a large number of proteins involved in chromatin organization and transcription initiation. This points to a putative role of IRF9 in altering the chromatin landscape and accessibility prior to interferon signalling.

Some of these interactors, such as BRD4 have previously been linked to interferon signalling^41,42^. Another novel IRF9 interactor, TRIM28, forms complexes with STAT1 and STAT3^43^. BioID with STAT2-BirA* identified several of the interactors identified in the IRF9 proximity screen. Human RVB1 and RVB2, which are highly conserved ATP binding proteins contained in chromatin-remodeling complexes such as Tip60/NuA4, interact with the transactivation domain of STAT2 in the nuclei of IFN-stimulated cells and are required for robust activation of IFNα-stimulated genes^44^. Although we could not find RVB1 or RVB2 among biotinylated proteins, Ep400, which is also a subunit of the Tip60/NuA4 complex, was shown in close proximity to both IRF9 and STAT2.

We conclude that complexes containing different ISGF3 subunits, both phosphorylated and unphosphorylated, serve distinct and relevant functions in controlling ISG expression. Our study focused on the STAT2-IRF9 to ISGF3 transition, but work by others addressed the importance of STAT1-STAT2 dimers. For example, their constitutive association may be relevant for the rapid formation of the tyrosine-phosphorylated heterodimer after IFN treatment^15^ and it prevents dissociation after dephosphorylation^45^. Furthermore, hemiphosphorylated pSTAT1-STAT2 dimers resulting from the preferential tyrosine phosphorylation of STAT1 dampen the response to IFNγ by restricting STAT1 access to the nucleus^18^. The multipurpose employment of ISGF3 subunits thus represents a striking example of the cell’s economy in the management of complex regulatory tasks.

## Acknowledgments

Funding was provided by the Austrian Science Fund (FWF) through grants P 25186-B22, SFB F6103 (to TD) and F6101 as well as F6106 (to MM). We thank M.Baccarini and G. Versteeg for critical reading of the manuscript and I. Steinparzer and P. Kovarik for advice concerning the RNA-seq and immunoprecipitation assays. We gratefully acknowledge discussion and suggestions by D. Levy and D. Panne that helped us to improve the manuscript. We also thank E. Ogris, S. Schüchner and F. Martys for their collaboration in the production of the monoclonal IRF9 antibody. Solexa Sequencing was performed by the VBCF NGS Unit (www.vbcf.ac.at). We thank A.Vogt, M.Farlik, C.Schmidl and C. Bücker for troubleshooting and suggestions with ChIP-seq experiments.

## Author contributions

E.P. designed and performed experiments, analyzed results and contributed to computational analyses of BioID data. D.D. and C.C. performed experiments and analyzed results A.S. performed experiments. M.H. and T.G. performed LC-MS and analyzed BioID protein interactors. M.M. provided intellectual input and expertise. M.N. performed computational analyses for RNA-seq and ChIP-seq. T.D. conceived the study, designed experiments, provided intellectual input, helped interpret data, provided supervision and secured funding. T.D. and E.P. wrote the manuscript.

## Materials & Correspondence

Correspondence and requests for materials should be addressed to T.D.

## METHODS

### Mice, animal experiments

Animal experiments were approved by the institutional ethics and animal welfare committee of the University of Veterinary Medicine, Vienna, and the national authority (Austrian Federal Ministry of Education, Science and Research) according to §§26ff of Animal Experiments Act (Tierversuchsgesetz TVG 2012, BGBl. I Nr 114/2012) under the permission license numbers BMWF 68.205/0032-WF/II/3b/2014 and BMWFW-68.205/0212-WF/V/3b/2016. Animal husbandry and experimentation was performed under the Austrian national law and the ethics committees of the University of Veterinary Medicine Vienna and according to the guidelines of FELASA which match those of ARRIVE. C57BL/6N, Irf9-/-, Stat1-/-, and Stat2-/- mice^1–3^ were backcrossed for more than 10 generations on a C57BL/6N background were housed in the same specific-pathogen-free (SPF) facility under identical conditions according to recommendations of the Federation of European Laboratory Animal Science Association and additionally monitored for being norovirus negative.

### Cell culture

Bone marrow-derived macrophages (BMDMs) were differentiated from bone marrow isolated from femurs and tibias of 8-to 12-week-old mice from both sexes. Femur and tibia were flushed with Dulbecco’s modified Eagle’s medium (DMEM) (Sigma-Aldrich) and cells were cultured in DMEM supplemented with 10% of fetal calf serum (FCS) (Sigma-Aldrich), 10% L929-cell conditioned medium as a source of colony-stimulating factor 1 (CSF-1), 100 units/ml penicillin, and 100 ng/ml streptomycin (Sigma-Aldrich). Cells were kept at 37°C and 5% CO_2_ and differentiated for 10 days. For ChIP-seq and RNA-seq experiments, BMDMs were differentiated in DMEM containing recombinant M-CSF (a kind gift from L. Ziegler-Heitbrock, Helmholtz Center, Munich, Germany). BMDMs and Raw 264.7 cells were stimulated with 10 ng/ml murine interferon gamma (IFNγ eBioscience; Cataog #14-8311-63) or 250 IU/mL of IFNβ (PBL Assay Science; Cataog #12400-1).

### Generation of monoclonal mouse IRF9 antibody

The murine monoclonal anti-IRF9 antibody was generated in collaboration with Egon Ogris, Stefan Schüchner and Florian Martys from the MFPL monoclonal antibody facility. Full length murine IRF9 was cloned into a pET-Duet1 (Novagen; Catalog # 71146) vector and expressed in E. coli Rosetta pLysS strain and purified using Ni-sepharose beads. Hybridomas from antibody-producing B cells and myeloma cells for the production of monoclonal IRF9 antibodies were generated. The best signal-to-noise ratio in ChIP and western blot analysis was obtained with the single clone 6F1-H5 which was used for this study. The purified antibody can now be purchased from Sigma (Anti-IRF-9, clone 6F1-H5, Cat. No. MABS1920, Emd Millipore).

### RNA isolation, cDNA synthesis and Q-PCR

Total RNA was extracted from mouse bone-marrow-derived macrophages using the NucleoSpin RNA II kit (Macherey-Nagel; Catalog #: 740955). The cDNAs was prepared using Oligo (dT18) Primer and the RevertAid Reverse Transcriptase (Thermo Fisher Scientific). Real-time qPCR experiments were run on the Mastercycler (Eppendorf) to amplify the Gapdh (housekeeping gene), using SybrGreen (Promega). Primers for qPCR and ChIP PCR listed in supplementary table 3.

### Western Blot

Cells were lysed in Laemmli buffer (120 mM Tris HCl pH 8, 2% SDS, 10% glycerol). Protein concentration was determined (Pierce BCA Protein Assay Kit). 30μg of protein were mixed with β-mercaptoethanol and bromophenol blue, boiled and loaded on a 10% SDS polyacrylamide gel.

Proteins were blotted on a PVDF membrane at 4 °C for 16 hours at 200 mA and for 2 hours at 400 mA in carbonate transfer buffer (3 mM Na2CO3, 10 mM NaHCO3, 20% Ethanol).

The membrane was blocked in 5% milk powder in TBS-T for 1 hour at room temperature. The membrane was washed three times with TBS-T and then incubated with the primary antibody over night at 4°C while shaking (Stat1 (Cell Signaling, Catalog #9172); Stat2 (Cell Signaling, Catalog #72604); Lamin B (Santa Cruz, Catalog #sc-6217); α-Tubulin (Sigma, Catalog #T9026); Phospho-Stat1 (Tyr701) (Cell Signaling, Catalog #9167); Phospho-STAT2 (Tyr689) (Cell Signaling Catalog #07-224); Lamin A/C (Santa Cruz, Catalog #sc-376248); GAPDH(Millipore, Catalog #ABS16); IRF9 (6F1)). The next day the membrane was washed three times again with TBS-T and incubated with an appropriate HRP-coupled secondary antibodies for 1h at room temperature (Jackson ImmunoResearch Inc. Code # 111-035-003, Code # 115-035-144). The membrane was analyzed with the ChemiDoc™ Imaging System from Bio-Rad.

### Nuclear and Cytoplasmic extraction

1 × 10^6^ bone marrow derived macrophages or Raw 264.7 cells were seeded in a 10 cm dish. The next day cells were treated for three hours with 500 nM staurosporine (Staurosporine solution from Streptomyces sp., Sigma S6942) or 15μM JAK Inhibitor (Pyridone 6, biovision, #2534) and afterwards stimulated for 30 minutes either with IFN-β or with IFN-γ. Extraction was carried out according to manufacturer’s instructions (NE-PER^™^ Nuclear and Cytoplasmic Extraction Reagents, Thermo Fischer, #78833). The whole nuclear fraction was loaded, whereas only 30% of the cytoplasmic fraction were loaded on a 10% SDS gel.

### Immunofluorescence

2 × 10^5^ BMDMs were seeded on glass cover slides, treated with IFN-β for 30 minutes and fixed with 3% paraformaldehyde for 20 minutes at room temperature. Cells were permeabilized with 0.1 % saponin in 0.5 M NaCl PBS. Blocking and all the stainings were carried out in 0.1 % saponin and 1 % BSA in 0.5 M NaCl PBS. The IRF9 antibody was used as a 1:10 dilution over night at 4 °C. Secondary goat anti-mouse Alexa Fluor^®^ 488 igGH+L (1:500; Catalog # A-11001) was purchased from Invitrogen. Samples were mounted in DAPI (ProLong^™^ Diamond Antifade Mountant with DAPI, Invitrogen, # P36962). Images were acquired using Zeiss Axio Imager Z2 with 63X oil objectives. Images were processed and analyzed using the ImageJ software. The background range in all images for DAPI was adjusted to 1500-16383 and for GFP 3000-16383. The images were changed to 14bit. A composite picture was made DAPI was set to to magenta and GFP to green. The type of picture was converted to RGB and a scale bar displaying 10 μm was inserted.

### RNA-seq

1.5 × 10^7^ cells were seeded on 15 cm dishes. The next day cells were stimulated for 2h either with IFN-β or with IFN-γ. 7 ml Qiazol Lysis Reagent (Qiagen) were added per 15 cm dish. Cells were scraped and vortexed for 20 seconds. 1 ml of the RNA samples in Trizol were used for the RNA prep and 200 μl of Chloroform were added. Samples were vortexed for 15 seconds and centrifuged for 5 minutes at full speed at room temperature. Supernatant was mixed with 1 volume Isopropanol, as well as 1/10 volume 5 M NaCl were added. Samples were incubated for 10 minutes at room temperature and centrifuged at 16.000g for 30 minutes at 4 °C. The pellets were washed twice with 75 % EtOH, dried and resuspended in 30 μl of dH_2_O.

For DNase treatment and clean up the RNase-free DNase Set (Qiagen, #79254) and RNeasy Mini Kit (Qiagen, #74104) were used. For library preparation, the NEBNext^®^ Poly(A) mRNA Magnetic Isolation Module together with the NEBNext Ultra II RNA Library Prep Kit from NEB #E7770L was used according to the manufacturer’s protocol. The samples were quality checked and sequenced at the Vienna Biocenter Core Facilities NGS Unit. The RNA-seq experiment was carried out as three independent biological replicates.

### Cloning

Full length STAT2 and IRF9 mouse cDNA were cloned into the pcDNA3.1 mycBioID vector (provided by Kyle Roux) and further subcloned into the pCW57.1 (Catalog # 41393) or pLVX-TRE3G-ZsGreen1 (Catalog #631350) and used for lentiviral transduction of Raw264.7 cells.

### BioID

BioID was performed according to a published protocol^4^. mycBioID was a gift from Kyle Roux (Addgene plasmid 35700).

5 × 10^6^ stable Raw 264.7 cells were seeded on 15 cm dishes and treated with 0,2 μg/ml doxycycline for 24h. 50 μM biotin were added for 18 additional hours. Cells were stimulated for 1.5h either with IFN-β or with IFN-γ. Cells were washed and lysed at room temperature (lysis Buffer: 50mMTris pH7.4; NaCL 500mM; 0,2% SDS; EDTA 5mM + 1x protease inhibitors). Triton X-100 and 50mMTris pH7.4 were added and the protein lysates were sonicated 2x for 30 seconds. Lysates were centrifuged for 5 minutes at full speed and supernatant was transferred to a new tube. Magnetic Pierce Streptavidin beads #88817 were washed 3x with lysis buffer. 105 μl beads were incubated with 1,3 mg of protein lysate over night at 4°C. 21 μl beads were kept for western blot analysis, the rest of the beads was used for the analysis with liquid chromatography mass spectrometry. Beads were washed at room temperature with wash buffer 1(2% SDS in H_2_O), wash buffer 2 (0,1% deoxycholic acid; 1% TritonX-100, 1mM EDTA, 500mM NaCl, 50mM HEPES; H_2_O) and wash buffer 3 (0,5% deoxycholic acid; 0,5% NP-40; 1mM EDTA; 250mM LiCl, 10mM Tris pH7.4;H_2_O). Beads were washed 5 times with 50mM Tris pH 7.4 and another 5 times with 50 mM ammonium bicarbonate (ABC) and then resuspended in 30 μL of 1 M urea in 50 mM ABC.

10 mM dithiothreitol was added and the samples incubated for 30 min at room temperature before adding 20 mM iodoacetamide and incubating for another 30 min at room temperature in the dark. Remaining iodoacetamide was quenched by adding 5 mM DTT and the proteins were digested with 300 ng trypsin (Trypsin Gold, Promega) at 37 °C overnight. After stopping the digest by addition of 1% trifluoroacetic acid (TFA), the supernatant was transferred to a new tube. The beads were washed with 30 μL 0.1% TFA, the supernatants were combined, and the peptides were desalted using C18 Stagetips^5^.

Peptides were separated on an Ultimate 3000 RSLC nano-flow chromatography system (Thermo-Fisher), using a pre-column for sample loading (Acclaim PepMap C18, 2 cm × 0.1 mm, 5 μm, Thermo-Fisher), and a C18 analytical column (Acclaim PepMap C18, 50 cm × 0.75 mm, 2 μm, Thermo-Fisher), applying a segmented linear gradient from 2% to 80% solvent B (80% acetonitrile, 0.1% formic acid; solvent A 0.1% formic acid) at a flow rate of 230 nL/min over 120 min. Eluting peptides were analyzed on a Q Exactive HF Orbitrap mass spectrometer (Thermo Fisher), which was coupled to the column with a nano-spray Flex ion-source (Thermo Fisher) using coated emitter tips (New Objective).

### Shotgun mass spectrometry data acquisition and processing

The mass spectrometer was operated in data-dependent mode, survey scans were obtained in a mass range of 380-1650 m/z with lock mass activated, at a resolution of 120k at 200 m/z and an AGC target value of 3E6. The 10 most intense ions were selected with an isolation width of 2 Da, fragmented in the HCD cell at 27% collision energy and the spectra recorded at a target value of 1E5 and a resolution of 30k. Peptides with a charge of +1 or > +6 were excluded from fragmentation, the peptide match feature was set to preferred, the exclude isotope feature was enabled, and selected precursors were dynamically excluded from repeated sampling for 30 seconds.

Raw data were processed using the MaxQuant software package (version 1.5.5.1;^6^ and the Uniprot mouse reference proteome (www.uniprot.org) as well as a database of most common contaminants. The search was performed with full trypsin specificity and a maximum of two missed cleavages at a protein and peptide spectrum match false discovery rate of 1%. Carbamidomethylation of cysteine residues were set as fixed, oxidation of methionine, phosphorylation of serine, threonine and tyrosine, acetylation of lysine, and N-terminal acetylation as variable modifications. For label-free quantification the “match between runs” feature and the LFQ function were activated^7^ - all other parameters were left at default.

MaxQuant search results were further processed using the Perseus software package (version 1.5.5.3,^8^. Contaminants, reverse hits, and proteins identified only by site were removed and the log2 transformed LFQ values were used for protein quantification. Mean LFQ intensities of biological replicate samples were calculated and missing values were replaced with a fixed value close to the detection limit. Only proteins that were quantified in both biological replicates in the samples of interest and displayed at least a 3-fold enrichment compared to the MYC-BirA control were considered to be enriched and represented using heat maps.

### Targeted mass spectrometry data acquisition and processing

PRM assays were generated based on the shotgun measurements, selecting up to 10 high intensity proteotypic peptides for STAT1, STAT2, and IRF9, with no missed cleavages, no methionine, and a good distribution over the chromatographic gradient. PRM assay generation was performed using Skyline^9^. After a test run with a pooled sample showing high target protein expression, we selected 6 or 7 peptides with a single charge state per protein according to their signal-to-noise ratio and their distribution over the gradient and designed a scheduled PRM assay with 6 min windows. For PRM data acquisition we operated the same instrument type as for shotgun MS, applying a 60 min gradient for separation and with the following MS parameters: survey scan with 30k resolution, AGC 1E6, 30 ms IT, over a range of 400 to 1300 m/z, PRM scan with 60 k resolution, AGC 1E5, 400 ms IT, isolation window of 1.2 m/z with 0.4 m/z offset, and NCE of 28%. In between samples, we acquired wash runs with the same method to monitor potential carry-over.

The data analysis, manual validation of all transitions (based on retention time, relative ion intensities, and mass accuracy), and relative quantification was performed in Skyline. Carry-over was negligible, all selected peptides eluted within the 6 minute time windows. The four to six most intense transitions were selected for each peptide and their peak areas were summed up for peptide quantification (total peak area). Peptide intensities were summed up to protein intensities. As quality control we extracted ion chromatograms (MS1) of eight peptides from three highly abundant background carboxylases which showed a LFQ ratio of roughly 1:1 in the shot-gun measurements and calculated normalization factors based on these peptides for the PRM dataset. Unnormalized and normalized protein ratios (log2) of IRF9, STAT1 and STAT2 were similar indicating good reproducibility over all measurements.

### Visualization of mass spectrometry data

The network diagram was generated with Cytoscape software^10^. Only proteins that displayed a three-fold enrichment in the samples of interest compared to the ligase only control were considered enriched and are displayed in the network. Spatial arrangement of nodes and colour code reflect grouping depending on whether proteins were already pre-associated in the absence of any signal or where specifically enriched upon 1.5h of IFN-β or IFN-γ. treatment. The ClueGO app^11^ was used to find overrepresented GO processes and a network of connected GO terms was created.

### ChIP and ChIP-seq

1,5 × 10^7^ bone marrow derived macrophages were seeded on a 15cm dish. The next day cells were stimulated for 1.5h either with IFN-β or with IFN-γ. Cells were crosslinked for 10 minutes at room temperature in 1% formaldehyde PBS (thermos fischer #28906). Cells were quenched with 0.125 M glycine for 10 min at RT. Cells were harvested and washed twice with ice cold PBS. Cells were centrifuged for 5 min at 1350 g at 4°C. Pellets were snap frozen in liquid nitrogen and stored at 80° C over night. Frozen pellets were thawn on ice for 60 minutes. Pellets were resuspended in 5 mL LB1 (50mM Hepes, 140mM NaCl, 1mM EDTA, 10% glycerol, 0,5% NP40, 0,25% TritonX100) by pipetting and rotated at 4°C for 10 min. Samples were centrifuged for 5 minutes at 1350 × g at 4 °C. Pellets were resuspend in 5 mL LB2 (10mM Tris, 200mM NaCl, 1mM EDTA, 0,5mM EGTA) by pipetting and rotated at 4 °C for 10 min. Samples were centrifuged for 5 minutes at 1350 × g at 4°C. Pellets were resuspend in 3 mL LB3 (10mM Tris, 100mM NaCl, 1mM EDTA, 0,5mM EGTA, 0,1% deoxycholate, 0,5% N-lauroylsarcosine). Samples were split into 2 × 1.5 mL in 15 mL polypropylene tubes suitable for the Bioruptor^®^ Pico (Diagenode). BioRuptor Sonicator settings: power = high, “on” interval = 30 seconds, “off” interval = 45 seconds, 6 cycles. Sonicated samples were centrifuged for 10 minutes at 16000 × g at 4 °C to pellet cellular debris. Chromatin concentration was measured by NanoDrop and 25 μg of chromatin were used for each IP. 300 μl 10% Triton X-100 were added to each 3 mL sonicated lysate. 25 μg of chromatin were stored at 4°C which served later on as an input control. Antibody of interest was added to sonicated chromatin aliquot and mixed (anti-STAT1 Santa Cruz sc-346; anti-STAT2 Santa Cruz Catalog # 07-140, IRF9 6F1-H5). All samples were filled up to 1ml with dilution buffer (16.5 mM Tris pH 8, 165 mM NaCl, 1.2 mM EDTA, 1% Triton X-100, 0.1% SDS, 0.1 mM PMSF, and cOmplete EDTA-free protease inhibitor cocktail (Sigma Aldrich)). Samples were rotated at 4 °C over night. 50 μl of magnetic beads (Dynabeads protein G, Life technologies, 10003D) per sample were blocked overnight in dilution buffer containing 1 % BSA at 4°C. The next day 50 μl of the beads were added to each sample and incubated at 4°C while rotating. Afterwards the beads were washed with 1 ml RIPA buffer (50 mM Tris HCL pH 8, 150 mM NaCl, 1% NP-40, 0.1% SDS, 0.5% sodium deoxycholate, 1 mM DTT), 2x High Salt buffer (50 Mm Tris pH 8, 500 mM NaCl, 0.1% SDS, 1% NP-40), 2x LiCl buffer (50 mM Tris pH 8, 250 mM LiCl, 0.5% sodium deoxycholate, 1% NP-40) and TE buffer (10 mM Tris pH 8, 1 mM EDTA) for 10 minutes at 4 °C. The samples were eluted in freshly prepared elution buffer (2% SDS, 100 mM NaHCO_3_, 10 mM DTT). The crosslink between proteins and DNA was reversed by adding 200 mM NaCl to each sample and incubation at 65 °C at 300 rpm for 12 hours. Proteinase K, 40 mM Tris pH 8 and 10 mM EDTA were added to each sample and incubated for 1 hour at 55 °C and 850 rpm. Each sample was transferred to a phase lock tube (5Prime), mixed 1:1 with phenol-chloroform-isoamylalcohol (PCI) and centrifuged for 5 minutes at 12000 × g. Supernatant was transferred and mixed with 800 μl 96% Ethanol, 40 μl 3M CH_3_COONa pH 5.3 and 1 μl Glycogen and stored for at overnight at −20 °C. Samples were centrifuged for 45 minutes at 4 °C and 16000 × g. Pellets were washed in ice cold 70% ethanol and dried at 65 °C, before diluting the DNA in H_2_O.

For library generation, the NEBNext Ultra II DNA Library Prep Kit for Illumina from NEB (Catalog #E7645S) was used according to the manufacturer’s protocol. The samples were quality checked and sequenced at the Vienna Biocenter Core Facilities NGS Unit.

### Immunoprecipitation

1.5 × 10^7^ bone marrow derived macrophages were stimulated for 1.5h with IFN-β. Cells were lysed in 1ml lysis buffer (10 mM Tris-HCl (pH 7.5), 50 mM NaCl, 30 mM NaPPi, 50 mM NaF, 2 mM EDTA, 1% Triton X-100, 1mM DTT, 0.1 mM PMSF and 1× protease inhibitor). Cells were incubated for 5 minutes on ice and then centrifuged for 5 minutes at 4°C, 12000 rpm. The supernatant was transferred to a new tube. 20 μl (10% of the lysate used for the IP) were used as an input control. 4 μg of an anti-STAT1 antibody (sc-346; Santa Cruz) or anti-STAT2 (catalog no. 07-140; Upstate) antibody, as well as 80μl of the IRF9 antibody were added to 200μl of lysate and incubated for 30 minutes at room temperature, while rotating.

50 μl of magnetic beads (Dynabeads protein G, Life technologies, 10003D) per sample were washed with Frackelton buffer. Then, 50 μl of the beads were added to each sample and incubated for 1h at room temperature while rotating. Afterwards the beads were washed three times with 1 ml frackelton buffer and proteins were eluted in SDS sample buffer (250 mM Tris-HCl pH 6.8, 20% Glycerol, 1.6% SDS, 20% β-Mercaptoethanol, 0.002% Bromophenol blue).

### Affinity pulldown of biotinylated ISRE probes

1.5 × 10^7^ Raw 264.7 macrophages were stimulated for 1.5h with IFN-β. Cells were harvested, washed and lysed in 1ml lysis buffer (10 mM Tris-HCl (pH 7.5), 50 mM NaCl, 30 mM NaPPi, 50 mM NaF, 2 mM EDTA, 1% Triton X-100, 1mM DTT, 0.1 mM PMSF, 1mM Na3VO4 and 1× protease inhibitor). Cells were incubated for 5 minutes on ice and then centrifuge for 5 minutes at 4 °C, 12000 rpm. 200 μl of each sample were incubated with 100 ng Oas1a (Fw Oas1a oligo 5’ bioteg 5’TAGATTTTCAGTTTCCATTTCCCGAGAAGGGCA 3’; Rv Oas1a oligo 5’TGCCCTTCTCGGGAAATGGAAACTGAAAATCTA 3’) and Isgf15 ISRE probes (Fw ISG15 oligo 5’ bioteg 5’ TATTTTCTGTTTCGGTTTCCTTTTCCTAC 3’; Rv ISG15 oligo 5’GTAGGAAAAGGAAACCGAAACAGAAAATA3’). The reaction was carried out in the presence of 0.1mM EGTA, 0,5mM DTT, 40mM KCL, 1mM MgCl2, 500ng competitor plasmid and 2μl Poly dI:dC (#20148E, Thermo Scientific). Samples were incubated at room temperature for 30 minutes while rotating. 50 μl of magnetic Pierce Streptavidin beads #88817 were added to each sample and incubated for 10 minutes at room temperature while rotating. Beads were washed three times with 1 ml binding buffer and proteins were eluted in SDS sample buffer (250 mM Tris-HCl pH 6.8, 20% Glycerol, 1.6% SDS, 20% β-Mercaptoethanol, 0.002% Bromophenol blue).

### Statistical analysis

mRNA expression data as well as ChIP data represent the mean values with standard error of mean (SEM). Differences in mRNA expression data or % of input data were compared using ratio t-test. All statistical analysis was performed using GraphPad Prism (Graphpad) software. Asterisks denote statistical significance as follows: ns, p>0.05; *, p≤ 0.05; **, p≤ 0.01; ***, p≤ 0.001. In all experiments n=3, where n represents the number of biological replicates.

### RNA-Seq Analysis

Reads mapping to mouse rRNA transcripts were removed using bwa/0.7.12 alignment^12^ and samtools/1.3.1^13,14^ Remaining reads were aligned to Mus musculus genome mm10 using TopHat v2.1.1^15^ (GTF annotation file mm10, RefSeq from UCSC, 2015/02). Reads in genes were counted with htseq-count v0.6.1.^16^ Differential expression analysis was carried out using DESeq 2 v1.16.1^17^, with an fdr threshold of 0.05.

Gene set enrichment analysis was performed using GSEA 3.0 against MsigDB v6.1 with gene abundance estimates in FPKM calculated using cufflinks v2.2.1.^18^

### ChIP-Seq Analysis

Raw reads were processed using the AQUAS TF pipeline (https://github.com/kundajelab/ChIP-Seq_pipeline;based off Encode phase-3;-idr_thresh 0.01), including alignment against the Mus musculus mm10 genome using BWA (v0.7.13^12^), deduplication using Picard MarkDuplicates (v1.126), Peak Calling using macs2 (v2.1.1^19^) and spp (v1.13^20^).

### Data availability

Raw and analyzed data reported in this paper are available under accession number GEO: GSE115435. Proteomics data will be uploaded to PRIDE and are available for reviewers from the corresponding author upon request.

